# Discrimination of melanoma cell lines with Fourier Transform Infrared (FTIR) spectroscopy

**DOI:** 10.1101/2020.09.05.284141

**Authors:** Bijay Ratna Shakya, Hanna-Riikka Teppo, Lassi Rieppo

## Abstract

Among skin cancers, melanoma is the lethal form and the leading cause of death in humans. Melanoma begins in melanocytes and is curable at early stages. Thus, early detection and evaluation of its metastatic potential are crucial for effective clinical intervention. Fourier transform infrared (FTIR) spectroscopy has gained considerable attention due to its versatility in detecting biochemical and biological features present in the samples. Changes in these features are used to differentiate between samples at different stages of the disease. Previously, FTIR spectroscopy has been mostly used to distinguish between healthy and diseased conditions. With this study, we aim to discriminate between different melanoma cell lines based on their FTIR spectra. Formalin-fixed paraffin embedded samples from three melanoma cell lines (IPC-298, SK-MEL-30 and COLO-800) were used. Statistically significant differences were observed in the prominent spectral bands of three cell lines along with shifts in peak positions. A partial least square discriminant analysis (PLS-DA) model built for the classification of three cell lines showed an overall accuracy of 92.6% with a sensitivity of 85%, 95.75%, 96.54%, and specificity of 97.80%, 92.14%, 98.64% for the differentiation of IPC-298, SK-MEL-30, and COLO-800, respectively. The results suggest that FTIR spectroscopy can differentiate between different melanoma cell lines and thus potentially characterize the metastatic potential of melanoma.

**Sources of Funding:** This research did not receive any specific grant from funding agencies in the public, commercial, or not-for-profit sectors.

## 1. Introduction

Melanoma is the most fatal type of skin cancer. Therefore, it is responsible for the majority of skin cancer-related deaths. Oncogenic transformation of melanocytes initiates the biological processes of melanoma followed by promotion and progression in the form of accumulative cellular damage, evasion of apoptosis, and epithelial-to-mesenchymal transition (EMT). This EMT process involves detachment from cell contact, alteration of cell shape, expression of matrix degrading enzymes, increased motility, and finally detachment from the primary tumour site and migration through the blood and lymphatic nodes, indicating the high metastatic potential that may lead to the development secondary tumours [1,2].

The diagnosis of melanoma is based on histomorphology and cytopathology of the tissue in hematoxylin-and-eosin (H&E) stained sections with adjunct histochemical and immunohistochemical staining. The stage or aggressiveness is then graded considering tumour thickness (Breslow index and Clark level), ulceration, mitotic rate, lymph node involvement and the distant metastasis status [3]. The concordance of histological diagnosis is only moderate in melanocytic lesions [4-6] and depends mostly on the experience of the pathologist. In addition to the morphological assessment of melanocytic tumours, Next-generation sequencing based analyses of the underlying genomic alterations are being developed and can be effective in the difficult intermediate category of melanocytic neoplasia [7].

Fourier Transform Infrared (FTIR) spectroscopy is based on the absorption of infrared light by a sample at the mid-infrared region. The biochemical information contained in the absorption spectrum can be used to probe the molecular structures and functions of measured samples. Compared with conventional histopathology, FTIR microscopy does not need any staining procedure prior to the measurement and can be performed directly on paraffin-embedded cells or tissues. Paraffin is often chemically dewaxed prior to the measurements. Alternatively, the contribution from paraffin can either be truncated [8] or digitally subtracted [9] from the acquired spectra. The acquired spectra analysis can be automated using machine learning algorithms, which may provide a label-free tool for automated objective classification of cells and extracellular components in the skin [10]. FTIR spectroscopy has previously been shown to discriminate melanoma from healthy skin and other skin lesions [11-14]. It has also been used to evaluate the effectiveness of drugs for treating melanoma [15]. Furthermore, Fourier Transform Infrared attenuated total reflectance (FTIR-ATR) modality has been used to assess the metastatic potential of live melanoma cells based on their membrane hydration level [16]. However, this procedure is not applicable to commonly used formalin-fixed paraffin-embedded (FFPE) biopsy samples. To our knowledge, no model has yet been developed to characterize the metastatic potential of fixed melanoma cells.

This study aims to develop a model that can discriminate between different melanoma cell lines based on their FTIR spectra. The cells used in the study are processed using the standard FFPE processing to make the results comparable with FFPE biopsy samples.

## 2. Materials and methods

### 2.1 Cell lines

Three melanoma cell lines, IPC-298 (ACC 251), SK-MEL-30 (ACC 151) and COLO-800 (ACC 193), were purchased from Leibniz-Institut DSMZ (Braunschweig, Germany) and used in this study. The melanoma cell line IPC-298 is derived from a primary melanoma tumour of a 64-year-old female, SK-MEL-30 is derived from a subcutaneous metastasis of a 67-year-old male with melanoma, and COLO-800 is derived from a subcutaneous nodule of a 14-year-old male with melanoma.

The cell lines were cultured in RPMI-1640 with 10% fetal bovine serum and 100 IU/ml penicillin and streptomycin (Pen-Strep solution HyClone laboratories, Inc. Utah, USA SV30010.01). The cells growing semi-confluent on a 6 cm cell culture dish were fixed with formalin for 30 minutes, and then the cells were scraped and centrifuged. The resulting cell pellet was mixed with 1.5% agar, followed by the paraffin infiltration (Tissue-Tek VIP 5 Jr, Sakura, The Netherlands) and embedding procedure. It took 13 hours to complete the paraffin-embedding process with a Tissue Tek processor. The cells were sequentially immersed in formalin, water, absolute alcohol, xylene and then infiltrated with molten paraffin in the processor. The infiltrated cells were then manually embedded into paraffin blocks. The detail of the paraffin infiltration and embedding procedure is shown in table 1.

**Table 1:** The sequential immersion of cells into various processing reagents within the Tissue Tek processor during the infiltration and embedding procedure of paraffin.

| **S.N.** | **Solutions** | **Time**<br>**(min)** | **Temperature**<br>**(°C)** | **Pressure vacuum**<br>**cycle (P/V)** | **Mix** |
| --- | --- | --- | --- | --- | --- |
| **1** | Formalin | 30 | 37 | On | Slow |
| **2** | Water | 30 | 37 | On | Fast |
| **3** | Ethanol (100%) | 30 | 37 | On | Slow |
| **4-8** | Ethanol (100%) | 60 | 37 | On | Slow |
| **9** | Xylene | 60 | 37 | On | Slow |
| **10** | Xylene | 90 | 37 | On | Slow |
| **11-12** | Paraffin<br>immersion | 60 | 57 | On | Off |
| **13** | Paraffin<br>embedding | 120 | 57 | On | Off |

Subsequently, 5-µm-thick sections were cut and placed on 2-mm-thick CaF_2_ windows for FTIR spectroscopic analysis. Paraffin was chemically dewaxed before the FTIR microscopy measurements. For dewaxing, the samples were immersed for 3 minutes in three different containers filled with xylene. To start rehydration, the samples were immersed for 3 minutes in three different containers filled with absolute alcohol, followed by immersion for 2 minutes in two containers filled with 96% ethanol. The samples were eventually washed with water to complete the rehydration process. Several 3-µm-thick sections of each cell line were also cut and placed on standard microscope slides and stained (after dewaxing) with haematoxylin and eosin using a standard staining protocol. The light microscope images of the haematoxylin and eosin-stained histological sections of the three melanoma cell lines are shown in figure 1.

**Figure 1:**
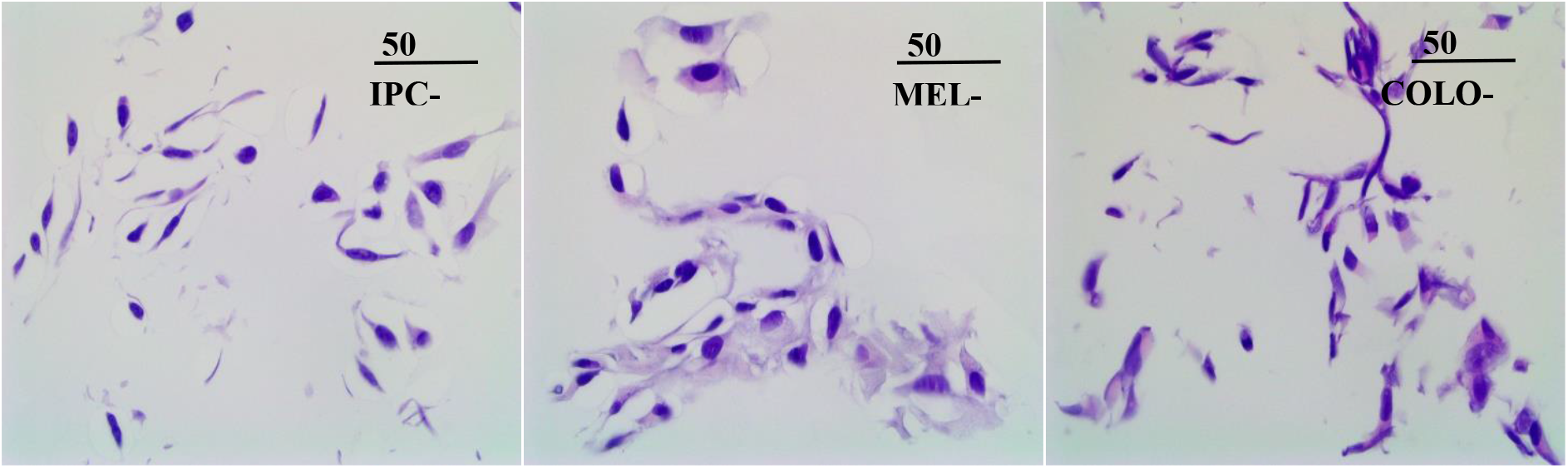
Histological images of the three melanoma cell lines used in the study stained with haematoxylin and eosin. (a) IPC-298 (b) SK-MEL-30 (c) COLO-800.

### 2.2 FTIR data acquisition

Nicolet iN10 MX (Thermo Scientific, Wisconsin, MA, USA) FTIR microscope was used to acquire spectra. A nitrogen-cooled mercury cadmium telluride (MCT) detector was used for the measurements. Two sections from each cell line were used in the study. A total of 260 cell spectra were acquired from the IPC-298 cell line (primary melanoma). In contrast, 300 and 260 cell spectra were acquired from SK-MEL-30 and COLO-800 cell lines (metastatic melanoma), respectively. Before each acquisition, a background spectrum was collected from a clean CaF_2_ window and was automatically subtracted from each cell spectrum. The spectra were collected in transmission mode, and one absorbance spectrum for each cell (aperture size: 20 ×20 µm^2^) was collected using the spectral range of 900-4000 cm^-1^ with a spectral resolution of 8 cm^-1^. The aperture size (20×20 µm) was selected to roughly match the cell size. This was done to minimize any signal outside the cells e.g. due to agar. It took about 4 minutes to acquire a spectrum of each cell with 256 scans (including background spectrum collection, which was also acquired with 256 scans).

### 2.3 Data pre-processing

For every spectrum, a simple offset correction for baseline was conducted prior to resonant Mie scattering correction (RMieS) to remove any negative values. After that, the RMieS algorithm [17] was applied to remove alterations due to the scattering of infrared radiation from the cells. The number of iterations was set to five, while the other correction options for the RMieS algorithm were set to the default values. After RMieS correction, spectra were normalized using vector normalization. The second derivatives were then calculated using the Savitzky-Golay algorithm (polynomial order = 2 and smoothing points = 7). Since the spectral features of paraffin remnants can often still be seen in the acquired spectra of deparaffinized samples [18], the lipid (2835-2999cm^-1^) and CH_3_ (1351-1480 cm^-1^) regions were truncated from further analysis as these regions are affected by paraffin [19]. In addition, the non-absorbing regions (1801-2834 cm^-1^ and 3631-4000 cm^-1^) were also removed from the spectra. The resulting second derivative spectra associated with five prominent spectral bands were used for the multivariate analysis. All processing was done in MATLAB (R2017b, MathWorks, Natick, MA, USA).

### 2.4 Data analysis

Firstly, the mean spectrum of each cell line was generated by averaging all spectra acquired from each cell line after required pre-processing procedures (baseline correction, RMieS correction and normalization) to visualize the spectral differences between the cell lines. The mean spectra consisted of multiple peaks associated with vibrational modes of molecules within the samples. The intensity and position of these peaks depend on the biochemical molecule (e.g., proteins, lipids and nucleic acid), their structure and conformation, and their intermolecular relationships [20]. The integrated areas of the peaks of interest were calculated to interpret the differences in the prominent spectral bands between the three cell lines, and their possible peak shifts were evaluated. Along with these peaks, the amide I/amide II ratio was calculated by dividing the integrated area of amide I by the integrated area of amide II. Furthermore, the one-way ANOVA test was performed, and the pairwise comparison between each cell line was conducted using the Bonferroni method to evaluate the significance of the differences of these bands between the cell lines. The limit of statistical significance was set at *p* < 0.05.

For the multivariate analysis, the second derivative spectra were calculated after RMieS correction. Subsequently, the spectra were truncated and normalized. In order to observe differences between the cell lines, Principal Component Analysis (PCA) was performed on the normalized second derivative spectra of the three cell lines. PCA does not use any *a priori* information about the classes of spectra but considers only the variance in the spectral data. PCA reduces a large set of variables to a small set of new variables while preserving most of the information. The new variables, i.e. principal components (PCs), are ranked by the amount of variance they explain: the first PC accounts for the largest variance in data while each succeeding PC accounts for the remaining variability. The result of PCA is commonly evaluated based on the scores of the first two or three PCs. In addition, the Partial Least Squares Discriminant Analysis (PLS-DA) was used to build a model for the classification of three melanoma cell lines. In this multiclass model, the spectra of each cell line formed their own class. Eight latent variables were used in the PLS-DA model. In the training phase, PLS-DA utilizes *a priori* knowledge about the classes of spectra of the training data set to determine the differences between the studied groups. A separate test set is then used to validate the trained model. PLS-DA model was built using 60% of the spectra from each cell line for training the model, while the remaining 40% were used as the test set. The model was run 50 times, and each time the spectra were randomly assigned to the training set and the test set.

## 3. Results

The FTIR spectra of the melanoma cell lines were investigated in the region 1000-4000 cm^-1^. The mean spectra (after required pre-processing) of the three malignant melanoma cell lines are presented in Figure 2. The spectra can be divided into three zones. In the first zone, i.e. the range of 1000-1480 cm^-1^, the absorption is mainly due to the cellular constituents of carbohydrates, nucleic acids and phosphates. The two prominent spectral bands in this region included in the analysis were observed at 1072 cm^-1^ and 1238 cm^-1^. The bands at 1072 cm^-1^ and 1238 cm^-1^ are symmetric and antisymmetric phosphate stretching vibration bands of nucleic acids and phospholipids. In the range of 1481-1800 cm^-1^, the main protein bands, i.e., amide II at 1539 cm^-1^ and amide I at 1655 cm^-1^, are observed. Amide I arise from C=O stretching vibration of peptide backbone and coupling of C-N vibration, while amide II is mainly due to C=O stretching vibration and coupling of N-H bending modes. The last zone, i.e., the range of 2835-3630 cm^-1^, results from the absorption of lipids and the N-H amino group. The only prominent band in this region included in the analysis was amide A observed at 3298 cm^-1^, which occurs due to the stretching vibration of NH groups of proteins. All of the cell lines well characterize the prominent spectral bands. [10]

**Figure 2:**
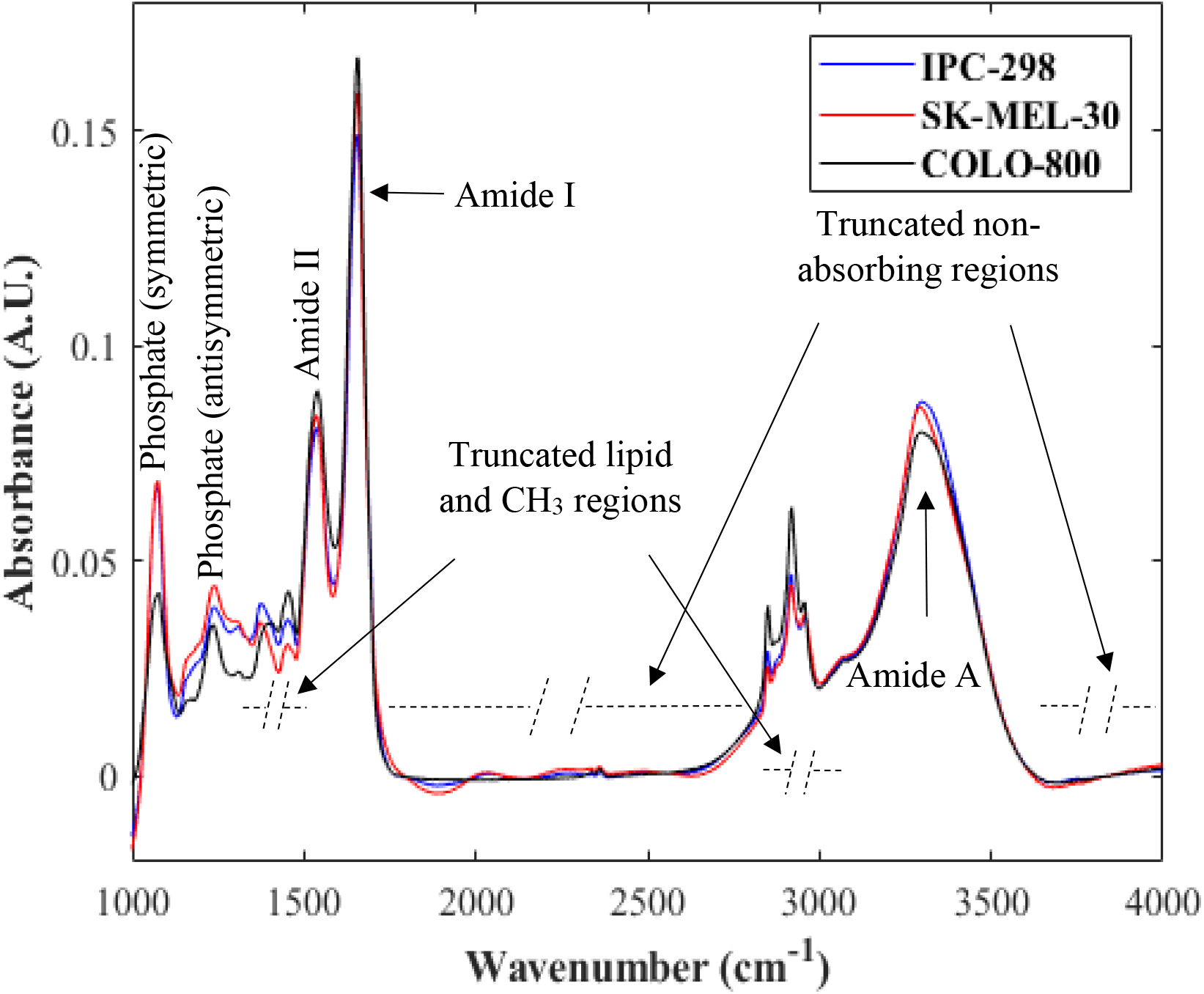
Normalized mean spectra of metastatic melanoma cells (SK-MEL-30: red and COLO-800: black) and primary melanoma cells (IPC-298: blue).

The spectra of the three melanoma cell lines show differences in integrated absorbances of prominent bands as well as band positions. Compared with the COLO-800 cell line, IPC-298 and SK-MEL-30 cell lines showed higher absorbance in symmetric phosphate, while the opposite was observed for antisymmetric phosphate. An increase in the absorbance intensities of amide I and amide II bands were observed in metastatic cell lines compared with primary, suggesting an alteration in native protein structure. The opposite was observed in the amide A region, where absorbance was weaker in the metastatic cell line. Compared with IPC-298, band shift to lower wavenumbers was observed in antisymmetric phosphate band of COLO-800 cell line (from 1238 cm^-1^ to 1234 cm^-1^), while in SK-MEL-30, amide II peak shifted from 1539 cm^-1^ to 1535 cm^-1^. Amide A band of both metastatic cell lines also shifted to lower wavenumber (from 3298 cm^-1^ to 3294 cm^-1^).

The phosphate and amide contents were calculated by measuring the integrated area of symmetric phosphate stretching (1000-1130 cm^-1^), antisymmetric phosphate stretching (1200-1273 cm^-1^), amide I (1585-1765 cm^-1^), amide II (1481-1581 cm^-1^) and amide A (3070-3630 cm^-1^) bands for all melanoma cell lines. Statistically significant differences were observed between each cell line in phosphate (antisymmetric), amide II and amide A bands. In phosphate (symmetric) and amide I bands, no significant differences were observed while comparing IPC-298 vs. SK-MEL-30 or IPC-298 vs. COLO-800 cell lines. However, the remaining comparisons showed significant differences in these bands as well. Moreover, the amide I/amide II ratio showed a significant difference between the primary and metastatic cell lines, but no significant difference was observed between the two metastatic cell lines. The peak positions of prominent spectral bands along with their integrated area are shown in Table 2. In contrast, the pairwise comparisons of each cell line are shown in Table 3.

**Table 2:** Wavenumbers of observed spectral bands with their integrated area (± SD) for statistical comparison.

| Peak assignment | Band Positions |  |  | Integrated area (mean) |  |  |
| --- | --- | --- | --- | --- | --- | --- |
|  | Primary (IPC-298) | Metastatic (SK-MEL-30) | Metastatic (COLO-800) | Primary (IPC-298) | Metastatic (SK-MEL-30) | Metastatic (COLO-800) |
| Phosphate (symmetric) | 1072 | 1072 | 1072 | $3.51 \pm 1.49$ | $3.63 \pm 1.40$ | $2.09 \pm 1.06$ |
| Phosphate (anti-symmetric) | 1238 | 1238 | 1234 | $0.34 \pm 0.15$ | $0.40 \pm 0.10$ | $0.49 \pm 0.11$ |
| Amide II | 1539 | 1535 | 1539 | $2.29 \pm 0.37$ | $2.63 \pm 0.37$ | $2.41 \pm 0.38$ |
| Amide I | 1655 | 1655 | 1655 | $6.87 \pm 0.50$ | $7.41 \pm 0.38$ | $6.85 \pm 0.52$ |
| Amide A | 3298 | 3294 | 3294 | $17.08 \pm 1.54$ | $16.63 \pm 1.23$ | $15.76 \pm 1.48$ |

**Table 3:**
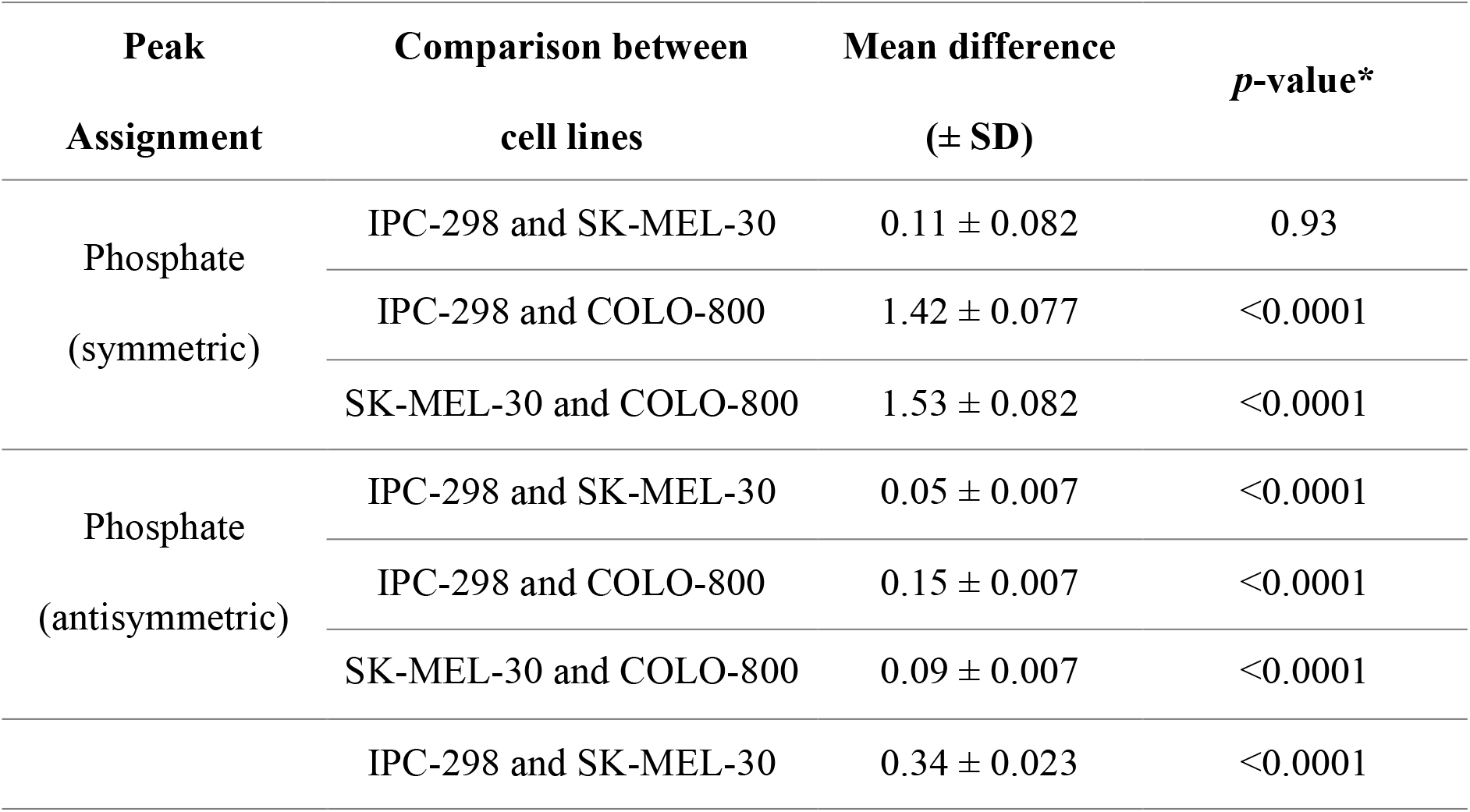
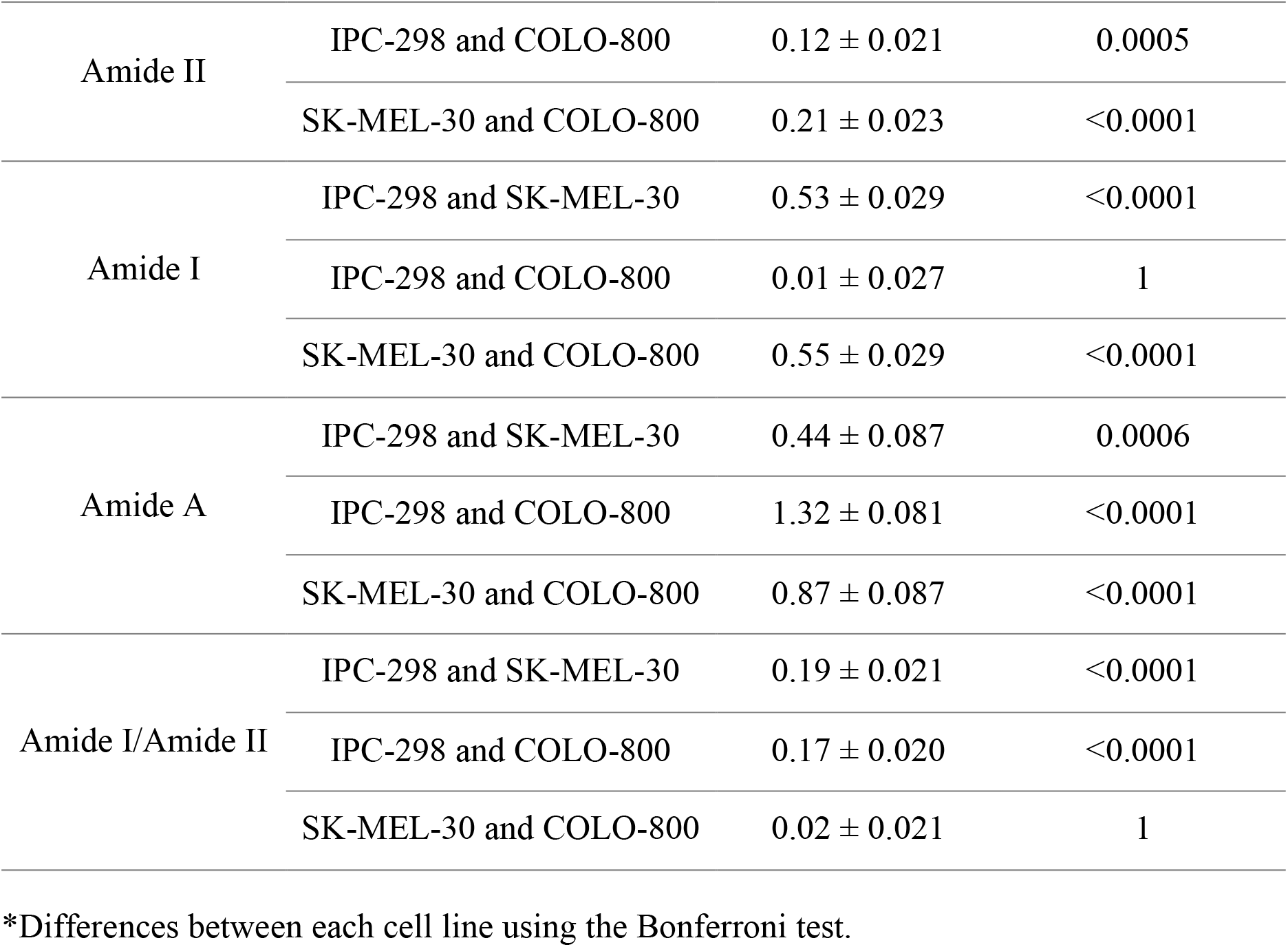
One-way ANOVA test for the prominent spectral bands between three malignant melanoma cell lines.

PCA was applied to all acquired spectra in the region 1000-4000 cm^-1^ (excluding paraffin affected and non-absorbing regions, i.e., 1351-1480 cm^-1^, 1801 - 2999 cm^-1^ and 3631-4000 cm^-1^). The first two PCs accounting for almost 57% of the variance (PC1: 33.63%, PC2: 23.23%) were used to explore the spectral differences in an unsupervised manner among the three cell lines. In Figure 3a, each point in the plot represents the transformed variable values of each spectrum known as scores. As seen in the figure, no definite clusters were formed and there was significant overlap between the three cell lines. Loadings of PC1 and PC2 were plotted for interpretation of the first two PCs (Figure 3b). In both PCs, contributions were observed in the carbohydrate and nucleic acid regions, which may be due to DNA and RNA structures, and in the amide I region, which are likely due to changes in protein secondary structures.

**Figure 3:**
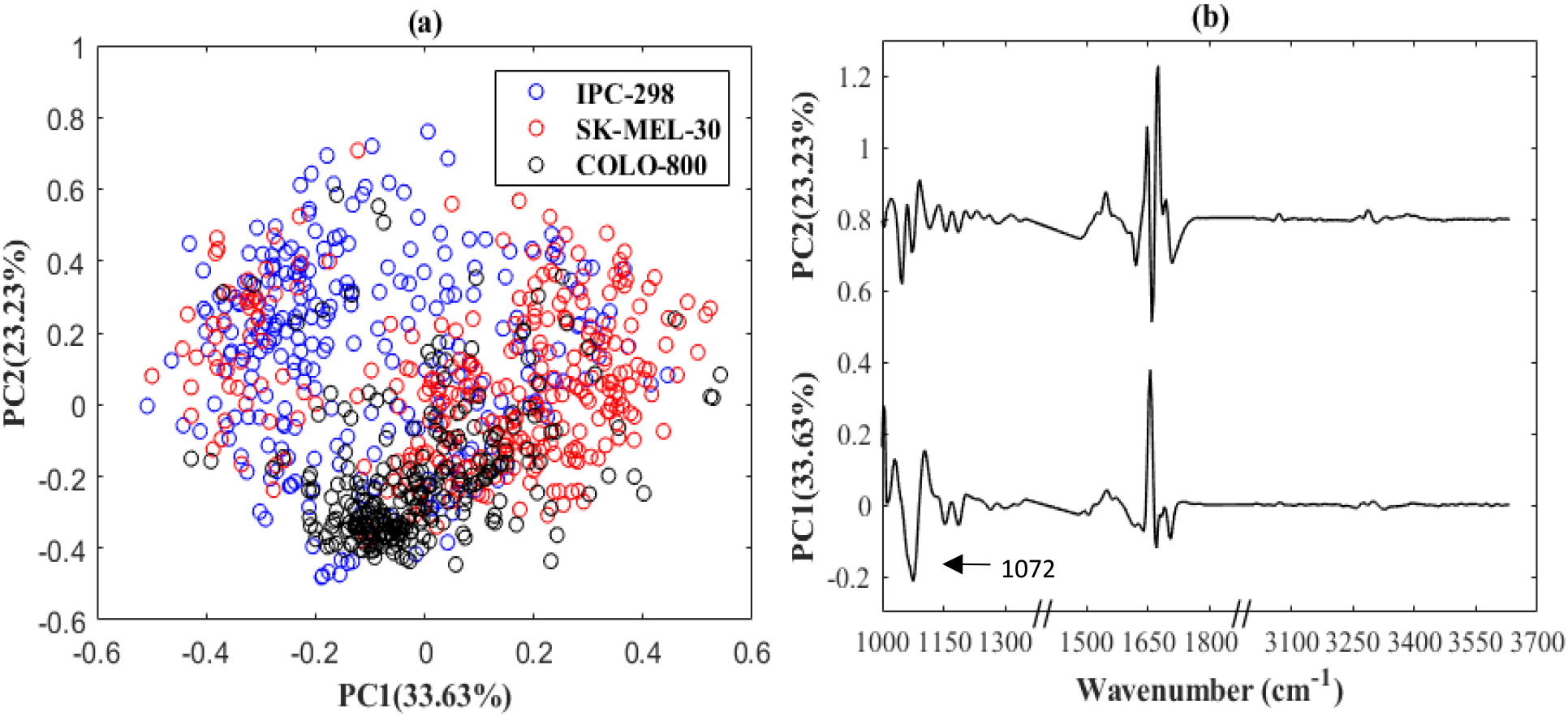
(a)Two-dimensional score plot computed using the first two PCs. The primary melanoma cells (IPC-298) are marked with blue circles and the metastatic melanoma cells: SK-MEL-30 and COLO-800 with red and black circles respectively. (b) The corresponding loading plots of the first two PCs. An offset value of 0.8 has been added to PC2 to avoid overlap.

To better classify the spectra of each cell line, PLS-DA model was created. The model showed high recognition rate with the mean sensitivities of 85%, 95.75%, 96.54% and the mean specificities of 97.80%, 92.14%, 98.64% for IPC-298, SK-MEL-30 and COLO-800, respectively. The predicted results for the test data set for each cell line are shown in tables 3. The predicted results are represented as the mean (± SD) percentages of the classification accuracy. The regression vectors obtained from the PLS-DA model for each cell line are shown in Figure 4. Large absolute regression coefficient values are observed in protein, DNA, and carbohydrate regions, indicating their importance for classification, while the high wavenumber region has only a small impact on classification.

**Table 3:** Recognition of each cell line with the PLS-DA model along with their sensitivity and specificity. The results are the average of 50 PLS-DA runs with random training/test set splits.

| Cell type | Predicted as |  |  | Sensitivity | Specificity |
| --- | --- | --- | --- | --- | --- |
|  | IPC-298 | SK-MEL-30 | COLO-800 |  |  |
| **IPC-298** | $85.00 \pm 3.75$ | $13.81 \pm 3.36$ | $1.19 \pm 1.28$ | $85.00 \pm 3.75$ | $97.80 \pm 1.02$ |
| **SK-MEL-30** | $2.75 \pm 1.44$ | $95.75 \pm 1.84$ | $1.50 \pm 1.02$ | $95.75 \pm 1.84$ | $92.14 \pm 1.76$ |
| **COLO-800** | $1.56 \pm 1.27$ | $1.90 \pm 1.22$ | $96.54 \pm 1.82$ | $96.54 \pm 1.82$ | $98.64 \pm 0.87$ |

**Figure 4:**
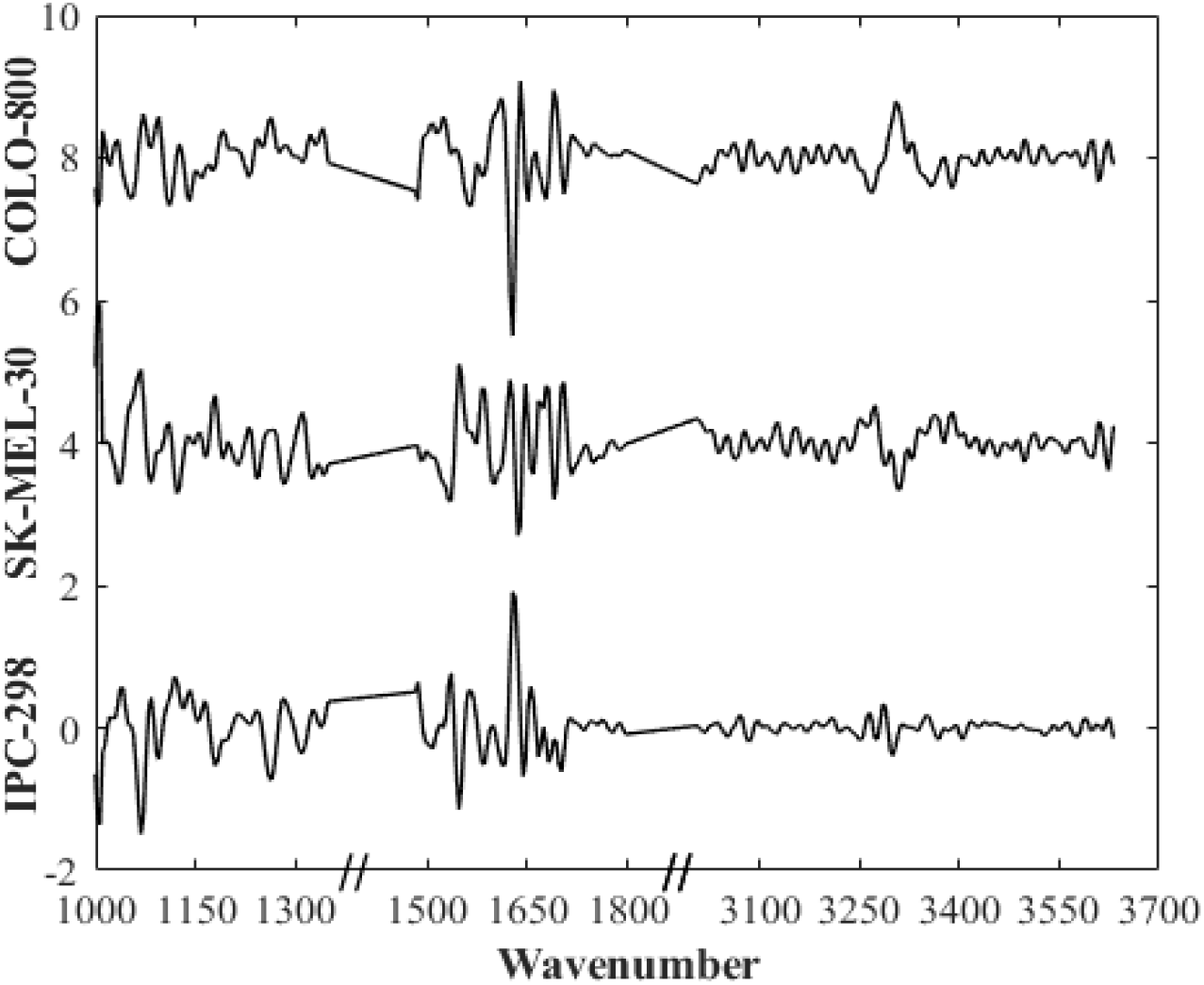
Regression vectors for the three classes used in PLS-DA model. Offset values of 4 and 8 were added to SK-MEL-30 and COLO-800 regression vector, respectively, to avoid overlap between the vectors.

## 4. Discussion

In this study, we evaluated the potential of FTIR spectroscopy for differentiating three melanoma cell lines processed using FFPE protocol. Firstly, the univariate analysis was used to show significant differences in integrated absorbances of phosphate (antisymmetric), phosphate (symmetric), amide I, amide II and amide A bands, between the FTIR spectra of each cell line. Then, multivariate analysis, i.e. PCA and PLS-DA, was performed for further discrimination. The developed PLS-DA models can accurately (>92%) differentiate each cell line from the remaining two cell lines. The result also suggests the possibility of differentiating primary melanoma cells from the metastatic melanoma cells with FTIR spectroscopy, but this requires further testing with larger sample sets.

Fixation of the cells is known to alter the cell FTIR spectra [21]. Furthermore, small but significant differences have been observed between air-dried and FFPE processed cell FTIR spectra [22]. In most cell studies, the samples are fixed to the substrate with air drying. However, virtually all morphology-based diagnoses conducted by pathologists are carried out on FFPE processed biopsies. We opted to use FFPE processing to ensure that the proposed approach is compatible with the standard protocols used for biopsy samples in pathology laboratories. The accurate classification of FFPE processed primary and metastatic melanoma cells is a promising sign for future studies where the metastatic potential of melanoma cells would be assessed from FFPE processed skin biopsies.

In our study, we observed high amide I and amide II absorbances for metastatic cells compared with primary melanoma cells. Furthermore, the amide II band shifted to a lower wavenumber in metastatic cells. This is likely a result of conformational alterations occurring in the secondary structures of proteins [23].

Another significant spectral variation was observed in phosphate bands. Asymmetrical phosphate band showed increased absorbance in metastatic cells. These phosphate stretching bands arise from the phosphodiester group of nucleic acids and provide information on various states of cells [24]. The changes in this band are due to alterations in the amount of nucleic acids, phospholipids and phosphorylated proteins produced in metastatic cells, which indicate an increased proliferative capacity of cells in metastatic condition.

Amide A region also showed significant spectral variation between the cell types. This band corresponds to N-H stretching vibration in proteins. The decrease in absorbance of this band is observed with the increasing metastatic potential of cells. Moreover, a considerable shift in the position of this band to a lower wavenumber was observed in metastatic cells. These changes are caused due to the increase of hydrogen bonding of the affected secondary amino function in metastatic cells [25]. These hydrogen bonds are essential in stabilizing the protein helix structure. The change in these structures indicates the alteration of the physiological environment.

PCA was carried out on all acquired malignant melanoma cell spectra. Successful separation of cell lines using this multivariate methodology would have been encouraging as it is an unsupervised method and does not need any prior knowledge for classification. However, the result from the analysis showed a limited separation with a significant overlap between the three cell lines. The similar phenotypic [26] and genomic [27] nature of the cells in the primary tumour site and the metastatic lesion may be the reason for this as mutational spectra are highly similar in matched primary and metastatic acral melanomas [28]. Furthermore, the phase of the cell cycle has been shown to produce significant alterations in the cell FTIR spectra [29,30]. Variations in the cell cycle may be another reason for the overlap between the cell lines in PCA.

A good classification was achieved for discriminating between different melanoma cell lines using PLS-DA. The developed model was able to predict each cell line with high sensitivity and specificity with the overall accuracy of 92.6%. In addition, the developed model also discriminated between primary and metastatic melanoma cells, suggesting the possibility of using FTIR spectroscopy with a developed model to characterize metastatic behaviour. Despite this promising result, it remains unclear which cell dependent and microenvironmental factors contribute the most to the classification. In a study by Wald *et al*. [26], melanoma cells in the primary tumour and metastasis were compared using FFPE tissue microarray sections. Neither supervised nor unsupervised analysis were able to reveal significant differences between the two classes. However, they were able to differentiate between batched primary tumours of patients at stage I or II (non-metastatic stages) and batched primary tumours of patients at stage III or IV (metastatic stages).

When a tumour develops, it undergoes a number of genomic alterations and becomes heterogeneous during the progression of the disease due to genetic, transcriptomic, epigenetic and phenotypic changes [31]. Genomic alterations differ among patients but also within the primary tumour and between the primary and metastatic lesion leading to intratumoral and intertumoral heterogeneity. Furthermore, metastatic cells can acquire new mutations and develop independently, known as intermetastatic heterogeneity [32]. These heterogeneities might explain the high level of irreproducibility of cancer research. The progression of melanoma into a metastatic pool of cells is greatly influenced by the tumour microenvironment, which is able to modulate various phenotypes reflecting therapy resistance [33]. Single-cell analysis has recently been recognized as the best approach to study intratumoral heterogeneity, but in contrast to FTIR spectroscopy, it would be difficult to implement it on a routine basis for diagnostics [34].

Although our study shows promising results in discriminating malignant melanoma cell lines, it has some limitations. Firstly, only three cell lines were included in the study and secondly, our samples consist of only cultured melanoma cells. Furthermore, the cell lines used in the study are from different patients without the same biological features and clinical evolution. In the future, a similar approach should be taken with biopsy samples, which are commonly used\ for histopathological inspection, encompassing of various other skin cells. This will reveal the real efficiency of the FTIR spectroscopic assessment of the metastatic potential of melanoma cells and tumour heterogeneity.

## 5. Conclusion

We observed significant differences between the FTIR spectra of different melanoma cell lines, and we were also able to discriminate between the cell lines with the developed PLS-DA model. The results encourage continuing the research towards the development of FTIR spectroscopic assessment of melanomas from skin biopsies.

